# Quantifying the risk of local Zika virus transmission in the continental US during the 2015-2016 ZIKV epidemic

**DOI:** 10.1101/298315

**Authors:** Kaiyuan Sun, Qian Zhang, Ana Pastore-Piontti, Matteo Chinazzi, Dina Mistry, Natalie E. Dean, Diana P. Rojas, Stefano Merler, Piero Poletti, Luca Rossi, M. Elizabeth Halloran, Ira M. Longini, Alessandro Vespignani

## Abstract

**Background:** Local mosquito-borne Zika virus (ZIKV) transmission has been reported in two counties of the continental United State (US), prompting the issuance of travel, prevention, and testing guidance across the continental US. Large uncertainty, however, surrounds the quantification of the actual risk of ZIKV introduction and autochthonous transmission across different areas of the US.

**Method:** We present a framework for the projection of ZIKV autochthonous transmission in the continental US during the 2015-2016 epidemic, using a data-driven stochastic and spatial epidemic model accounting for seasonal, environmental and detailed population data. The model generates an ensemble of travel-related case counts and simulate their potential to trigger local transmission at individual level.

**Results:** We estimate the risk of ZIKV introduction and local transmission at the county level and at the 0.025° x 0.025° cell level across the continental US. We provide a risk measure based on the probability of observing local transmission in a specific location during a ZIKV epidemic modeled after the one observed during the years 2015-2016. The high spatial and temporal resolutions of the model allow us to generate statistical estimates of the number of ZIKV introductions leading to local transmission in each location. We find that the risk is spatially heterogeneously distributed and concentrated in a few specific areas that account for less than 1% of the continental US population. Locations in Texas and Florida that have actually experienced local ZIKV transmission are among the places at highest risk according to our results. We also provide an analysis of the key determinants for local transmission, and identify the key introduction routes and their contributions to ZIKV spread in the continental US.

**Conclusions:** This framework provides quantitative risk estimates, fully captures the stochasticity of ZIKV introduction events, and is not biased by the under-ascertainment of cases due to asymptomatic infections. It provides general information on key risk determinants and data with potential uses in defining public health recommendations and guidance about ZIKV risk in the US.

## Background

From 2015 to 2016, the Zika virus (ZIKV) epidemic spread across most countries in the Americas, including the United States (US) [1–3]. As of September 17, 2017, three US territories, including Puerto Rico, have reported 37,009 ZIKV cases mostly due to widespread local transmission [3, 4]. Laboratory evidence of possible ZIKV infections has been found in 4,341 pregnant women from US territories, 139 of whom have had pregnancy outcomes with ZIKV-related birth defects [3, 5, 6]. The continental US and District of Columbia have reported 5,464 travel associated ZIKV cases, including 2,155 pregnant women with evidence of ZIKV infection and 103 ZIKV-related birth defects [3]. Two geographical locations have experienced local transmission of ZIKV in the continental US: Miami-Dade County, in Florida and Cameron County, in Texas [7, 8]. While the epidemics in Florida and Texas were limited, the indirect impact on the local economy has been remarkable [9].

Concerns have been raised that several other locations of the continental US are at risk of ZIKV transmission, thus triggering a number of studies aimed at identifying populations at highest risk of local transmission [10–20]. In particular, detailed studies based on environmental suitability, epidemiological factors and travel-related case importations have been used to estimated the risk for specific counties in the US [21, 22].

In this study, we quantify the risk of local ZIKV transmission by using a data-driven stochastic and spatial epidemic model accounting for seasonal, environmental and population data. The model also accounts for the association of socioeconomic status and the risk of exposure to mosquitoes, and it has been previously used to estimate the introduction of Zika in the Americas and the spatial and temporal dynamics of the epidemic [23]. By using an extensive likelihood analysis with data from places with a reliable epidemiological surveillance system, the model generates a stochastic ensemble of simulations estimating the place and time of introduction of ZIKV in Brazil and the unfolding of the epidemic in the Americas. For each simulation, the individual-level scale of the model allows the construction of daily travel-related case counts (TCC) in the continental US at the county-level and at the spatial resolution of 0.025° × 0.025° (approximately *2.5km x 2.5km)* cells, comparable in size to the ZIKV active transmission areas identified by the Centers for Disease Control and prevention (CDC) in Florida [24]. Using the time series of TCC and the mechanistic transmission model, it is possible to estimate the probability, that a specific location will experience local ZIKV transmission during the 2015-2016 time window. The methodology proposed here provides a statistical estimate of ZIKV transmission risk that is not biased by the under-ascertainment of infections and the single historical occurrence of the case importation timeline that fail to account for the full stochasticity of transmission events. The synthetic ZIKV travel-related infection data also allows us to identify key sources and routes of ZIKV introductions. Results from our study can provide guidance for public health agencies in their efforts to identify populations and seasons at high risk of ZIKV transmission, so that resources towards outbreak prevention and response can be allocated more efficiently.

## Methods

We consider three major factors associated with local ZIKV transmission in the continental US: the intensity of travel-related infection importations, the environmental suitability for ZIKV transmission, and the population socioeconomic status. In this study, we develop a data-driven computational framework (Figure 1) to quantitatively account for these three factors and to evaluate their impact on ZIKV transmission. Based on this framework, we assess the risk of autochthonous ZIKV transmission across the continental US through the full course of the 2015-2016 ZIKV epidemic.

**Figure 1:**
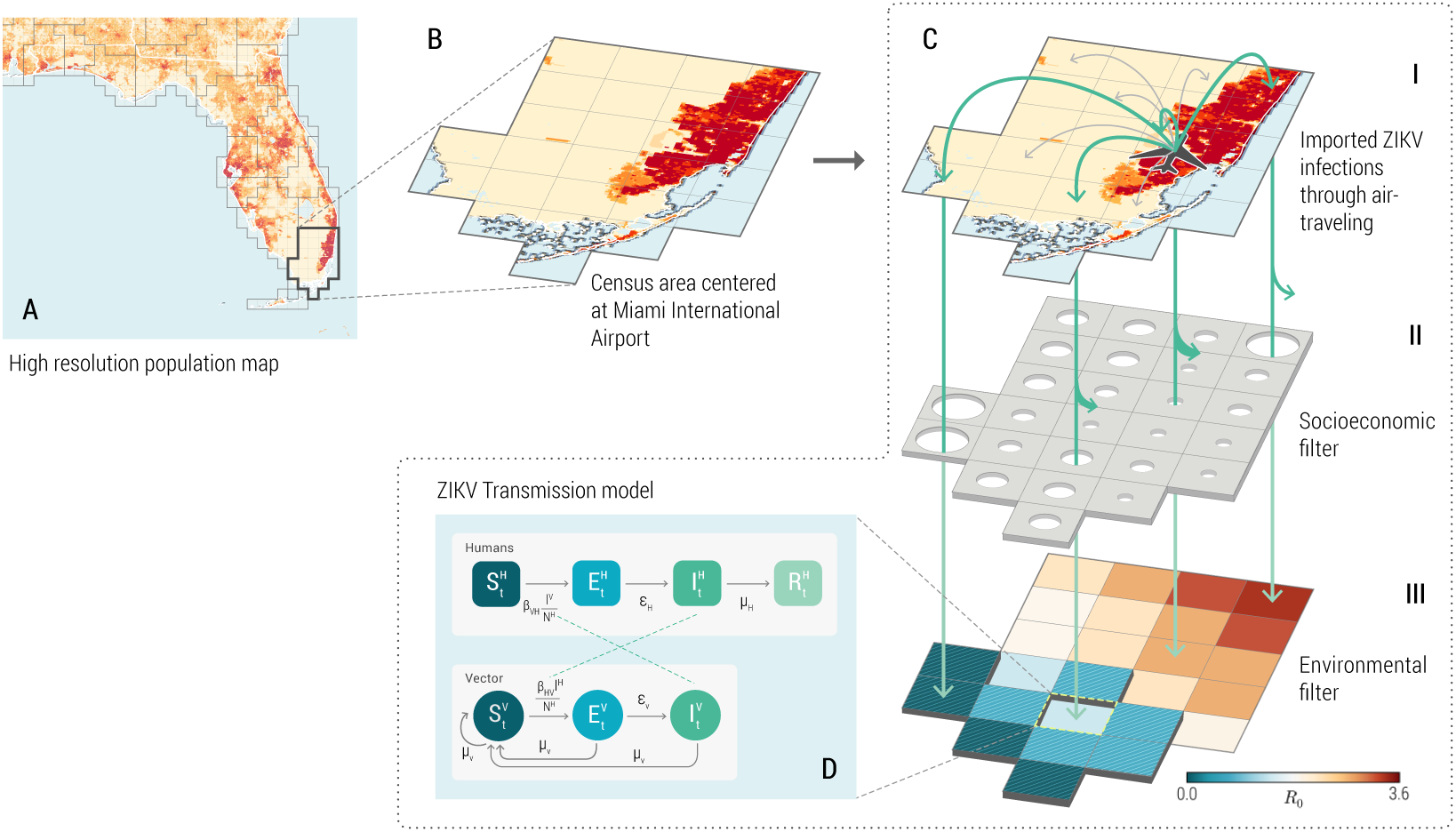
A schematic illustration of the computational framework to assess the risk of ZIKV introductions into the continental US. **(A)** High resolution (0.025° × 0.025° ~ 2.5*km* × 2.5*km*) population density map [48] and Voronoi tessellation of the continental US into census areas with a major airport transportation hub at each of their centers [49]. **(B)** An example of the census area centered at Miami International Airport. **(C) I:** Travel-associated ZIKV infections entering the Miami International Airport. Location of residence of each ZIKV infection is randomly assigned with likelihood proportional to the population density within each census area. **II:** The probabilistic filter of the risk of exposure to mosquitoes due to socioeconomic factors such as housing conditions, sanitation, and disease awareness. **III**: Spatiotemporal specific ZIKV transmission dynamics are influenced by environmental factors that are temperature sensitive, including the spatial distribution of *Aedes* mosquitoes, seasonal mosquito abundance, and ZIKV transmissibility. **(D)** Compartmental stochastic ZIKV transmission model used to evaluate the environmental suitability of ZIKV transmission. Humans are divided into susceptible *S^H^*, exposed *E^H^*, infectious *I*^*H*^, and recovered *R*^*H*^ compartments, and mosquitoes are divided into susceptible *S*^*V*^, exposed *E*^*V*^ and infectious *I*^*V*^ compartments.

The starting point of our methodology is the construction of a synthetic database of TCC entering the US through airport transportation. The database is generated from simulations based on a large-scale spatial model simulating the 2015-2016 ZIKV epidemics, where both symptomatic and asymptomatic ZIKV infections are considered [23]. The synthetic database of TCC contains for each infected individual the time of arrival, stage of ZIKV infection, airports of origin and arrival, and location of residence in the continental US. A schematic sample of the database is shown in Table 1.

**Table 1:** A sample of the database containing simulated travel-related ZIKV infected individuals entering the US.

| case ID | time of arrival | airport of arrival | stage of infection | airport of origin | location of residence (latitude, longitude) |
| --- | --- | --- | --- | --- | --- |
| 0001 | 2015-12-01 | MIA | Exposed | BOG | (25.864, -80.257) |
| 0101 | 2016-07-15 | JFK | Infectious | SJU | (40.729, -73.991) |
| 0212 | 2016-11-23 | MIA | Infectious | SJU | (25.808, -80.130) |

Each infected individual’s likelihood of exposure to mosquito bites and his/her capability of triggering autochthonous ZIKV transmission is not only determined by the ecological presence of mosquitoes in his/her location of residence [25, 26]; the individual’s socioeconomic status, which is strongly associated with factors such as sanitation conditions, accessibility to air conditioning, level of disease awareness, also affects the likelihood of exposure to mosquitoes [14, 27, 28]. Our computational framework considers a data layer based on global socioeconomic indicators [29], which is calibrated with historical mosquito-borne disease outbreaks in naive populations to provide a likelihood map of the individuals’ exposure to mosquitoes [23]. This map serves as a spatial filter (Figure 1 C-II) that probabilistically selects individuals exposed to mosquito bites down to the resolution of a 0.25° × 0.25° cell containing his/her location of residence. Each of the exposed individuals can potentially trigger detectable local ZIKV transmissions (Figure 1 C-III, D), according to the stochastic mechanistic ZIKV transmission model that takes into account mosquito abundance, the current temperature in the area, and the transmission dynamics of ZIKV (see Appendix file 1: Supplementary Information). Due to the fine spatial and temporal resolution, the transmission model is able to account for the significant variability in the ZIKV basic reproduction number (*R*_0_) across locations, as well as the variability within the same location at different times. These differences in *R*_0_ are driven by temperature and the mosquito abundance, among other variables. We define a detectable local transmission as the generation of 20 or more autochthonous transmission infections triggered by a single ZIKV infection introduction. Smaller outbreaks will likely go unnoticed assuming a 5% to 10% detection rate of infections due to the large proportion of asymptomatic cases [30–32]. The details of the mechanistic model and the calculation of the socioeconomic likelihood of exposure are reported in the Appendix file 1. More technically we can define the following procedure:

1. We randomly sample one out of the simulated TCC from the statistical ensemble output of the ZIKV model [23].
2. For each infected individual in the TCC, we stochastically determine whether he/she is potentially exposed to mosquito bites based on the probability of exposure *p*_*e*_ at the location of residence *x*. *p*_*e*_ is calibrated based on socioeconomic indicators and *x* identifies a specific county or spatial cell. In each location *x*, these individuals could potentially trigger local transmission.
3. Based on the individual’s stage of infection (exposed or infectious), time of introduction, and location of residence (at 0.025° × 0.025° resolution, comparable to the sizes of ZIKV active transmission areas in Florida designated by the CDC [3].), we simulate local ZIKV transmission with the stochastic transmission model used in the global model [23] (described in Additional file 1: Supplementary Information) and other studies [33, 34] with the specific parameters calibrated to each 0.25° × 0.25° cell in the US. We consider that an imported infection is triggering a detectable outbreak, if he/she generates a chain of more than 20 infections acquired through autochthonous transmission.
4. For each simulated TCC, the above procedure identifies all the infections triggering detectable local transmission. For every time interval Δ*t* and geographical area *x* of interest, we can associate the variable *n*(*x,* Δ*t*) = 1 if there is at least one imported infection from the TCC that triggers detectable local transmission, and *n*(*x*, Δ*t*) = 0 otherwise.

In order to provide a probabilistic risk measure, we execute *N* = 10^6^ resamplings from the ensemble of simulated TCC generated by the model and repeat the above procedure. The resampling procedure accounts for the many possible TCC compatible with the observed ZIKV epidemic and stochastic effects in the local transmission. This is because not all case importations will result in local outbreaks, even in areas where transmission is favored. The risk of local ZIKV transmission for area *x* during time window Δ*t* can be thus defined as

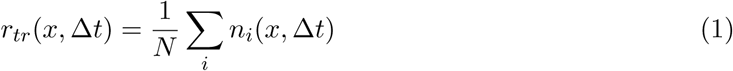

where *i* indexes the 10^6^ outcomes from the resampled TCCs. This definition of the risk can be aggregated at various spatial (≥ 0.025° × 0.025°) and temporal resolutions (≥ 1 *day*), and it can be used to generate risk maps of ZIKV introduction across the continental US. Unless otherwise specified, we consider in this manuscript the local transmission risk *r*_*tr*_(*x*) that is defined on the Δ*t* referring to the time window spanning from January 1, 2015 to December 31, 2016. This definition of risk can be interpreted as the probability of observing local transmission in a specific area per ZIKV epidemic. If an area has a risk of 0.1, we can expect to observe local transmission in that area in one out of ten equivalent ZIKV epidemics. This definition is akin to the one defined for other hazards such as flooding.

## Results

By using the methodologies outlined in the previous section, we provide quantitative estimates of *r*_*tr*_(*x*) at both the county and 0.025° × 0.025° cell resolution. Figure 2 (A) shows the risk of ZIKV introduction at county level in the continental US through the full course of the simulated 2015-2016 ZIKV epidemic. We consider four main brackets for the risk and the associated population sizes. At the county level, the largest risk bracket *r*_*tr*_(*x*) > 0.5 includes only 0.71% of the total population in the continental US. In these areas one would expect local transmission during every other major ZIKV epidemic equivalent to the one that swept through the Americas in 2015-2016. Only 2.56% of the continental US population has a risk of local transmission larger than in 1 out of every 8 epidemics similar to the one of 2015-16, i.e. *r*_*tr*_(*x*) > 0.125. Thus the risk of local transmission is extremely concentrated in specific geographical locations. Figure 2 (D) shows the population living in counties with different risk brackets of ZIKV introduction and their percentage with respect to the total population in the continental US. In many cases, risk is measured over time. In this case, one should factor in the time of recurrence of a ZIKV epidemic with magnitude similar to the one observed in 2015-2016. For instance, a previous study suggests the next wave for a large scale ZIKV epidemic may not return within 10 years due to herd immunity effects [35]. According to the study, a 1/16 risk would translate into a 160 *years* recurrence interval.

**Figure 2:**
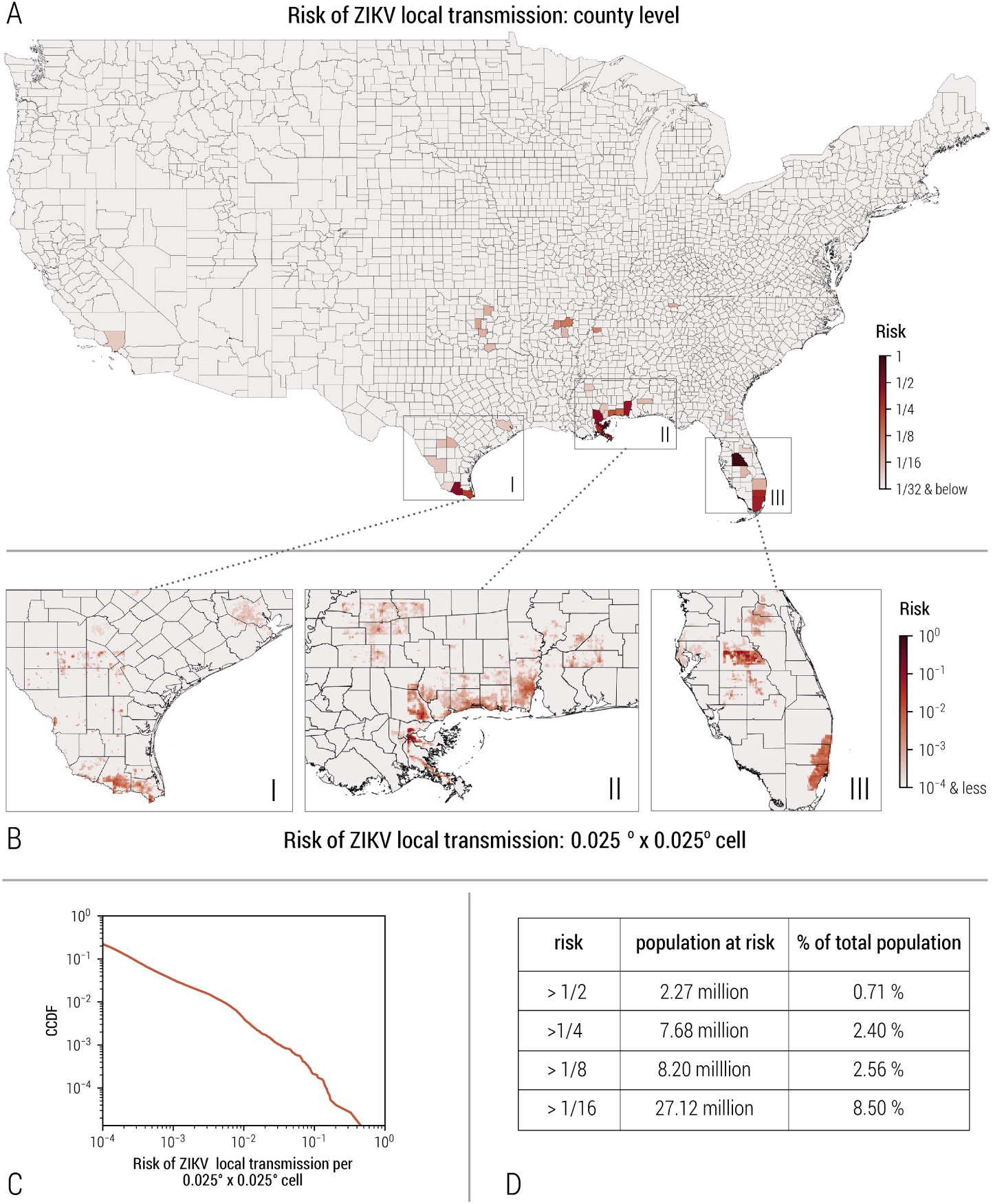
The cumulative risk of local ZIKV transmission in the continental US. The cumulative risk of local ZIKV transmission at different spatial resolutions are evaluated through the full course of the simulated 2015-2016 ZIKV epidemic. **(A)** The cumulative risk map of local ZIKV transmission for each county in the continental US. The color scale indicates for any given county the probability of experiencing at least one ZIKV outbreak with more than 20 infections (details in Additional File 1). **(B)** High spatial resolution estimates (0.025° × 0.025°) of the cumulative risk of local ZIKV transmission through the full course of the simulated 2015-2016 ZIKV epidemic. **(C)** The complementary cumulative distribution function of the local ZIKV transmission risks for all 0.025° × 0.025° cells (on a log-log scale). The heavy tail feature of the distribution reflects strong spatial heterogeneity in terms of local ZIKV transmission risk. **(D)** The total population in the counties of the US with different risk levels of local ZIKV transmission and their percentage with respect to the total population in the continental US.

The counties of Miami-Dade, Florida, and Cameron, Texas, where local transmission was observed in the year 2016, are both estimated to be high-risk locations (risk bracket: greater than 1/4). Other densely populated areas along the Gulf Coast also show up as high-risk locations, in agreement with estimates from other models [12]. The risk of ZIKV introduction and local transmission *r*_*tr*_(*x*) is spatially very heterogeneous (Figure 2 (A), (B)). This heterogeneity persists even within the state of Florida, where most areas are estimated to be environmentally suitable for ZIKV transmission all year long [12, 36]. This is mostly because of socioeconomic and local climate heterogeneities. At a spatial granularity of 0.025° × 0.025°, it is possible to perform a statistical analysis of the risk distribution. In Figure 2 (C) we report the probability distribution that any cell has a given *r*_*tr*_ (*x*). The distribution has a very right-skewed heavy tail extending over more than four orders of magnitude; a clear signature of the large heterogeneity of the risk in the continental US.

It is worth stressing that the source of ZIKV introductions in each location is time-dependent, since the TCC is determined by both the magnitude of the epidemic, and travel patterns in the regions in the Americas affected by ZIKV. Our model explicitly simulates travelers with ZIKV infections at the individual level, with detailed information about the traveller’s origin and destination at the daily scale. This allows us to decompose the relative contribution of potential ZIKV introductions from different epidemic regions and to identify routes of high risk with high spatiotemporal resolution. In Table 2, we report the likelihood of local ZIKV transmission in Miami-Dade, Florida, for the year 2015 and 2016 imported from the Caribbean, Central America, and South America. The likelihood accounts for intensity of ZIKV transmission in epidemic regions, the travel volume between the source regions and Miami-Dade, as well as the the time-dependent environmental suitability of local transmission in Miami-Dade. In Figure 3, we report the daily risk of ZIKV infections in Miami-Dade from different geographical regions as well as the time-dependent relative contributions of different regions to the risk throughout the years 2015 and 2016.

**Table 2:**
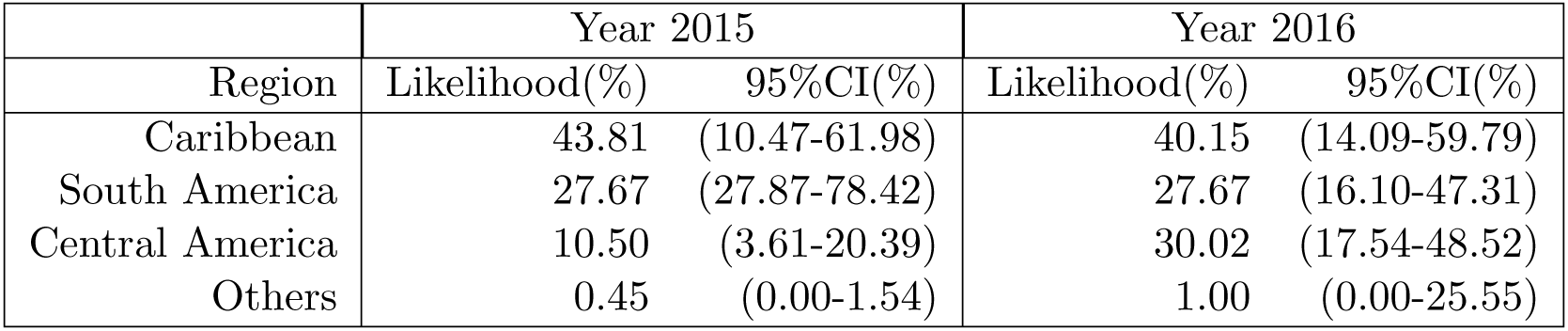
The likelihood of a given local ZIKV transmission event in Miami-Dade, Florida caused by ZIKV importation coming from different geographical regions (Caribbean, South America, Central America, Others) for the years 2015 and 2016.

**Figure 3:**
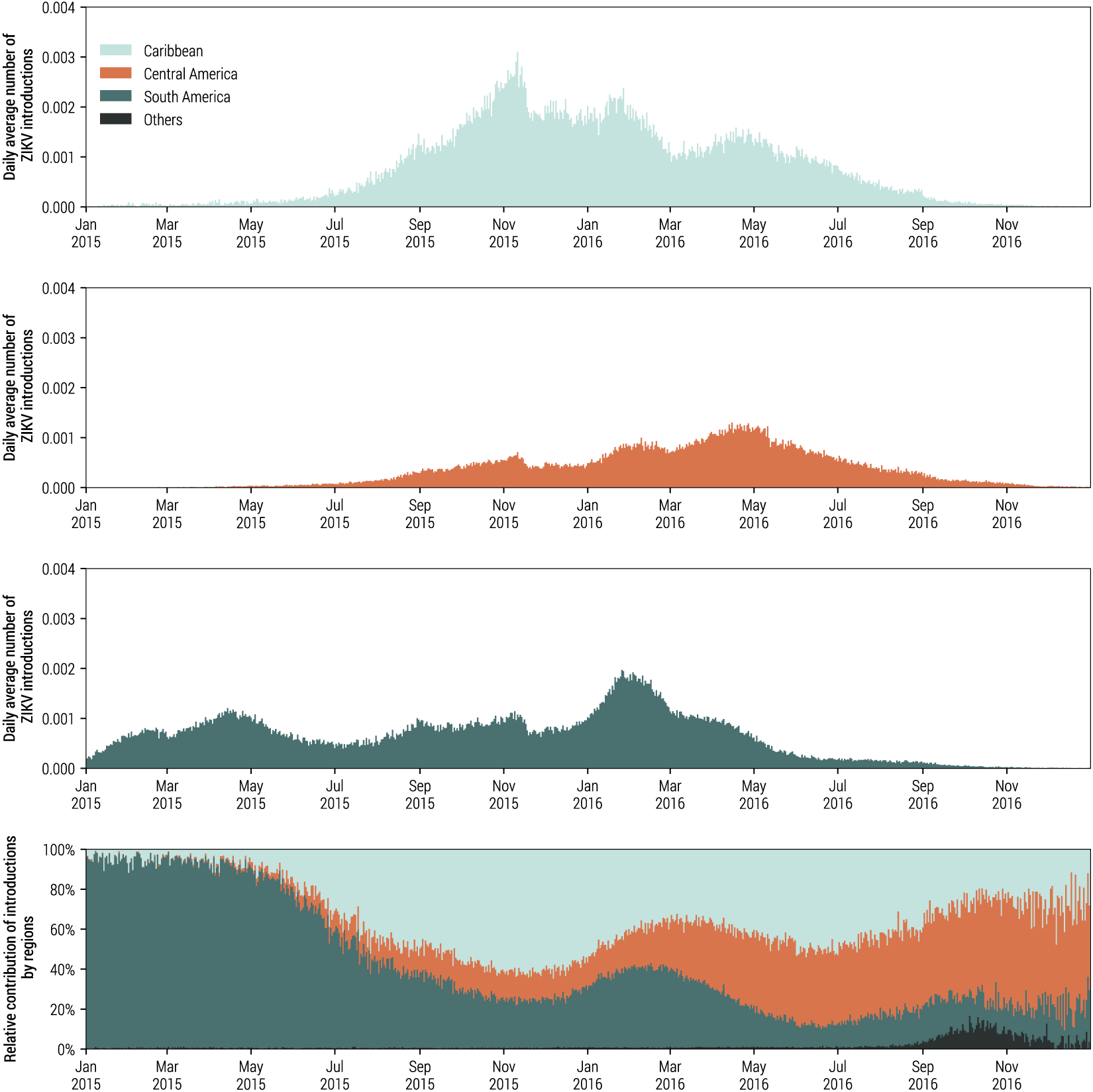
A breakdown of local ZIKV transmission events by the geographical origins of travel-associated ZIKV infections in Miami-Dade, Florida. Row 1-3 display the daily average number of imported infections per day that trigger outbreaks with more than 20 infections, originating from the Caribbean, Central America, and South America. Row 4 displays the relative contributions to the expected number of local ZIKV transmission events by different geographical regions.

As shown in both Table 2 and Figure 3, in 2015, countries in the Caribbean and South America are the major contributors to ZIKV introduction risk in Miami-Dade. On the other hand, countries in Central America become the major contributors in 2016. This reflects the fact that ZIKV epidemic started earlier in South American countries, including Brazil and Colombia, and later on spread to countries in Central America. Caribbean countries, however, remain a major source of infection importation in both 2015 and 2016. This is possibly due to the high travel volumes between Florida and the Caribbean, as well as high incidence rate and weak seasonality of ZIKV transmission in that region. This is in line with epidemiological data from Florida’s Department of Health, as well as phylogenetic analysis based on sequenced ZIKV genomes from both infected humans and mosquitoes in Florida [37].

In Figure 4, we zoom in on three representative areas to disentangle the key determinants shaping the spatiotemporal risk of local ZIKV transmission. Panel A, B and C in Figure 4 represent geographical areas covering Miami-Dade, Florida, Cameron, Texas, and New York City, New York. Both Miami-Dade and New York City have experienced a high volume of ZIKV infection importations due to high population density and close proximity to major international transportation hubs. Cameron, Texas, on the other hand, had far fewer ZIKV infection importations. However, due to socioeconomic factors, the population in Cameron, Texas, is more likely to be exposed to mosquitoes than the populations of Miami-Dade and New York City. Consequently, the volume of Cameron’s imported infections that are exposed to mosquito bites is comparable to those of Miami-Dade and New York City.

**Figure 4:**
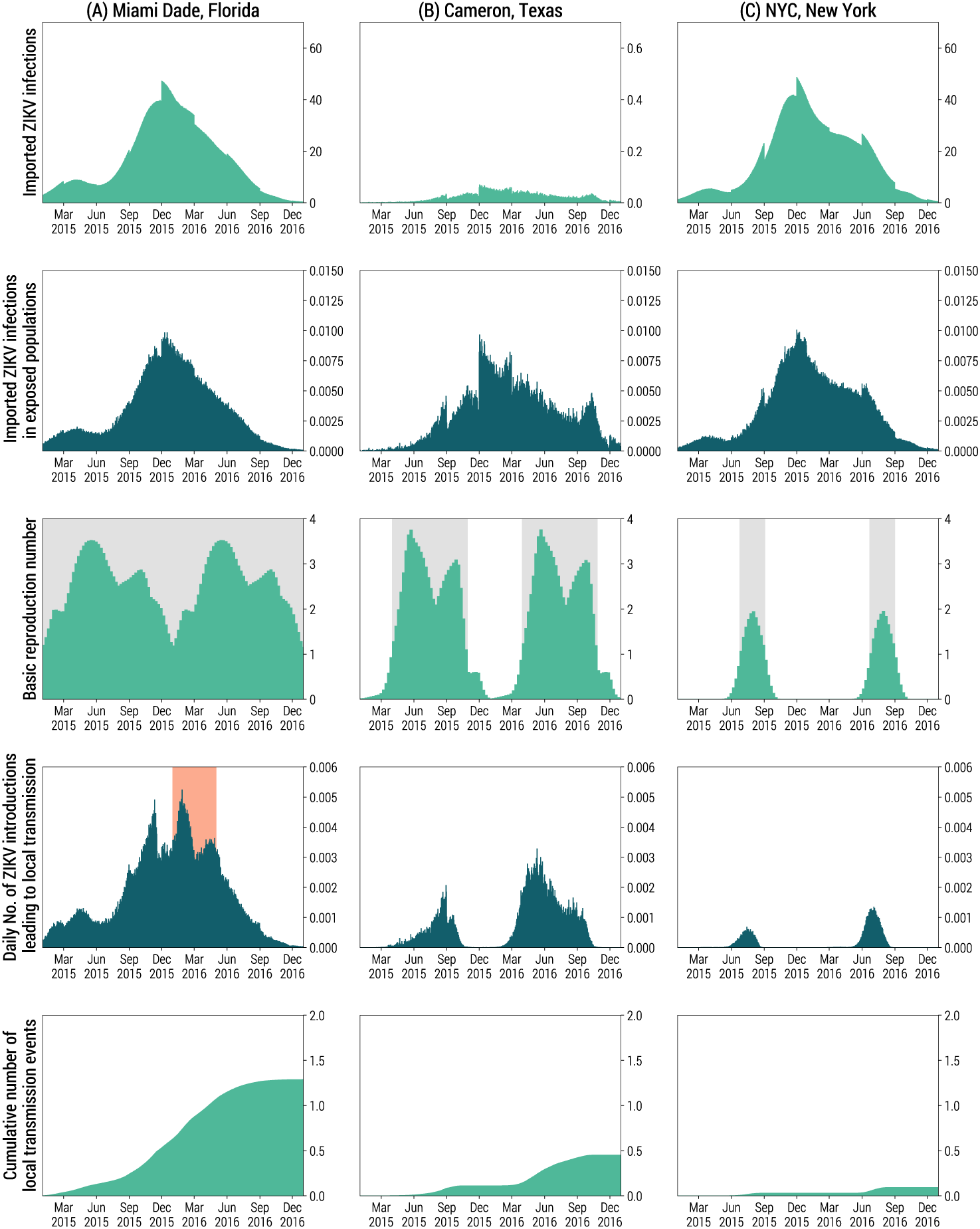
Factors which co-shape the spatiotemporal risk of local ZIKV transmission in three different regions in the continental US. Columns from left to right represent: **(A)** Miami-Dade, Florida, **(B)** Cameron, Texas, and **(C)** New York City, New York. Three major factors contribute to the spatiotemporal risk of local ZIKV transmission: the intensity of ZIKV importation, the socioeconomic risk of exposure to mosquitoes, and the environmental suitability of ZIKV transmission. Row **(1)** shows the average daily number of imported ZIKV infections. Note that for Cameron, Texas, the scale on the y-axis is different than that of Miami-Dade, Florida, and NYC, New York. Row **(2)** shows the average number of imported ZIKV infections that pass through the socioeconomic filter *p*_*e*_ and reside in areas potentially exposed to mosquitoes. Row **3** shows the basic reproduction number (weekly average) calculated based on the ZIKV transmission model. Grey shaded time windows indicate when the basic reproduction number *R*_0_ > 1 and sustainable ZIKV transmission is possible. Row **4** shows the expected daily number of ZIKV introductions with the red shaded time window indicating the estimated time of local ZIKV transmission based on phylogenetic analysis [37]. Row **5** shows the average cumulative number of local ZIKV transmission events since January 1, 2015.

The environmental suitabilitiy of ZIKV transmission in the three areas however is remarkably different. The basic reproduction number *R*_0_ is above the epidemic threshold (*R*_0_ > 1) in Miami-Dade throughout the year, indicating ZIKV transmission is environmentally suitable all year long. Cameron, Texas, has moderate environmental suitability, where *R*_0_ drops below the threshold in winter seasons. New York City is far less suitable for ZIKV transmission environmentally, with a narrow time window of approximately two months during summer when *R*_0_ is larger than one.

Given the individual-level resolution of the model, we can focus on the daily average number of individual travel infections leading to local transmission. This is a different indicator than risk. The latter is defined as the probability of observing at least one event of detectable local transmission in the area, thus overlooking the number of different introduction events that trigger local transmission. The profile of daily ZIKV introductions that will lead to local transmission (Figure 4, row 4) is jointly shaped by ZIKV infection importations, socioeconomic risk of exposure, as well as the environmental suitability of ZIKV transmission. The cumulative number of ZIKV introductions leading to local transmission is high in both Miami-Dade, Florida, and Cameron, Texas, where local transmission occurred in the year 2016. The time of ZIKV introduction in Miami-Dade, Florida is estimated to have occurred between January and May 2016 based on phylogenetic analysis of sequenced ZIKV genomes from infected patients and *Aedes aegypti* mosquitoes [37]. Our model suggests (Figure 4, row 4) high risk of ZIKV introduction during the same time window, despite relatively low environmental suitability. The high risk of introduction in Miami-Dade between January and May 2016 is mainly driven by a high influx of imported ZIKV infections. Based on our simulations, Miami-Dade county has on average 1.29 cumulative introductions leading to local transmission events (95%CI [0-9]) throughout 2015 and 2016 (Figure 4, row 5, insert). However, the distribution of the number of introductions is positively skewed (skewness *γ*_1_ = 4.40), with a maximum of 55 introductions. This indicates the possibility of multiple introductions during the ZIKV outbreak in Miami-Dade, Florida, in line with estimates from phylogenetic analysis [37].

In order to investigate to what extent the spatial variation of local ZIKV transmission is driven by key socioeconomic and environmental determinants, we consider a regression model exploring the relation between the average number of local ZIKV transmissions (log(*n*_*tr*_) is the dependent variable) and three key determinants: the number of ZIKV importations, average temperature and the GDP per capita. Specifically, the explanatory variables include:

1. log(*N*_*im*_), the logarithm of the cumulative average number of TCC for each 0.25° × 0.25° cell from January 1, 2015, to December 31, 2016.
2. log(*f*_20°_), the logarithm of the fraction of days over the year with an average temperature larger than 20° Celsius for each 0.25° × 0.25° cell. [38].
3. log(*GDP*), the Gross Domestic Product per capita [29, 39], in terms of purchasing power parity [23] for each 0.25^°^ × 0.25^°^ cell.

In Table 3, we show that if all three explanatory variables are included in the regression (Model 1), the model can explain 73.9% of the variance in the number of average introductions leading to local transmission in each cell x. While only considering log(*N*_*im*_) and log(*f*_20°_) (Model 2) we can explain 56.2% of the variance, and using log(*N*_*im*_) (Model 3) alone can explain 47.5% of the variance. It is worth remarking that such a simple statistical analysis cannot fully explain the variance of log(*n*_*tr*_) due to the nonlinear dependency between ZIKV transmission, vector population dynamics, and temperature. It is also due to the highly nonlinear nature of the disease transmission dynamics captured by the epidemic threshold (where the basic reproduction number (*R*_0_) needs to be larger than one to be able to spread in a population). In addition, more than 90% of the geographical areas in the continental US are not included in the regression because the simulations project no local transmission events in those areas. Indeed, 77% (in terms of areas) of these “risk-free” areas are not environmentally suitable for ZIKV transmission according to our model.

**Table 3:** Regression analysis between log(*n_tr_*) and explanatory variables including log(*N*_*im*_), log(*f*_20°_) and log(*GDP*). Specifically, denotes the average number of local ZIKV transmissions within each 0.25° × 0.25° cell from January 1, 2015 to December 31, 2016; *N*_*im*_ denotes the number of ZIKV importations; *f*_20°_ denotes the fraction of days with temperature higher than 20 degree Celsius, GDP denotes the Gross domestic product per capita in purchasing power parity. In Model 1, all three explanatory variables log(*N*_*im*_), log(*f*_20°_) and log(*GDP*) are included. Model 2 includes log(*N*_*im*_), log(*f*_20°_). Model 3 only includes log(*N*_*im*_). For each model we report the regression coefficient (95%CI) for each of the explanatory variables along with R squared and the number of observations.

|  | Model 1 |  | Model 2 |  | Model 3 |  |
| --- | --- | --- | --- | --- | --- | --- |
| $\log(N_{im})$ | 0.72*** | (0.69, 0.74) | 0.48*** | (0.45, 0.51) | 0.49*** | (0.46, 0.52) |
| $\log(f_{20^\circ})$ | 2.18*** | (1.89, 2.48) | 3.40*** | (3.01, 3.78) | | |
| $\log(GDP)$ | -13.29*** | (-14.20, -12.38) | | | | |
| R squared | 74.9% |  | 57.9% |  | 47.5% |  |
| Observations | 1220 |  | 1220 |  | 1220 |  |

\*\*\*p<0.01

## Discussion

A prominent feature of our findings is the spatiotemporal heterogeneity of ZIKV transmission risk across the continental US. Spatially, our model estimates that approximately 68.9% of the people in the continental US live in areas that are environmentally suitable for ZIKV transmission, in line with other models’ estimates [40]. However, taking all ZIKV introduction and transmission determinants into consideration, areas with non-negligible risk (greater than 1/8) are concentrated in densely populated areas along the Gulf Coast, capturing 2.56% of the US population. From a temporal perspective, certain areas experience strong seasonality of ZIKV environmental suitability, with a narrow time-window when ZIKV transmission is possible. Given limited resources, identifying seasons and regions of high risk may help guide resource allocation for high risk population screening, intervention, and vector control.

Our model is also able to identify the high-risk routes of ZIKV importations through air travel. Imported infections originating from Caribbean countries serve as a major contributor to trigger local ZIKV transmission in Florida. Though it has the highest number of estimated ZIKV infections among all countries, Brazil is not a major contributor overall (5.75% of potential introductions leading to local transmission across the continental US). This is due to Rio de Janeiro and Sao Paulo, two of the largest transportation hubs in Brazil which make up 65% of the international travels to US from Brazil, being located in the Southern region where ZIKV transmission activity is relatively low. In addition, Rio de Janeiro and Sao Paulo have the opposite seasonality compared to the continental US. When ZIKV transmission is environmentally suitable in Rio de Janeiro and Sao Paulo, ZIKV transmission is not suitable in most of the continental US. Thus imported ZIKV infections from Brazil are less likely to fuel potential transmissions in the US.

Our model also suggests that in Miami-Dade, Florida, the overall risk of ZIKV introduction in 2015 is comparable to that in 2016, although local transmission was only observed in 2016. This could be explained just by the stochasticity of transmission events. Another possibility is that because of the high asymptomatic rate of ZIKV infections, limited local transmission events occurred in 2015 without being picked up by the surveillance system. Awareness of ZIKV was low in 2015 as the World Health Organization declared ZIKV as a Public Health Emergency of International Concerns only in early 2016. Around the same time the CDC announced a Health Alert Network advisory for Zika virus [3], marking the start of active monitoring of ZIKV activities in the US.

The proposed model has several limitations. The high volume of cruise ship stops along coastal areas of Florida to the Caribbean may elevate the risk of ZIKV transmissions beyond what is estimated in our model. Sexual transmission and transmission through other routes, not considered by our model, may facilitate the risk of local transmission even further. From January 1, 2015, to August 9 2017, there were 49 reported ZIKV cases in the continental US acquired through other routes, including sexual transmission [3, 41–43]. This indicates that a larger population may be affected by ZIKV [44–46]. In addition, ZIKV RNA was detected in semen as long as 92 days after symptom onset and is able to be sexually transmitted 31-42 days after symptom onset [47]. ZIKV’s ability to persist in infected males and the potential to infect through sexual transmission long after symptom onset is troublesome. In areas where strong seasonality of ZIKV transmission is presented, ZIKV could linger in an infected individual to pass through seasonal barriers and trigger local transmission when environmentally suitable. However, the specific risk through sexual transmission or other transmission routes is not well understood, and the overall impact of ZIKV infections acquired through other routes remains unclear. As such we do not include them in our study. Risk of exposure to mosquitoes associated with socioeconomic factors is widely recognized but poorly quantified. In our model we utilize seroprevalance studies from nine chikungunya outbreaks on confined, naive populations to estimate this association, in line with other approaches used to estimate the ZIKV attack rate [14]. Further studies however are needed to advance our understanding of the association between risk of exposure to mosquitoes and socioeconomic status.

In 2017-2018, ZIKV transmission activities in most countries throughout Latin America has plummeted [2], in agreement with models’ estimates [23, 35]. The risk of ZIKV introduction in the continental US is expected to be negligible as imported infections triggering the local transmission would be drastically reduced. However, one should exercise caution as vector transmitted diseases are known to show strong spatial heterogeneity and seasonality, and are affected by socioeconomic factors. The stochastic nature of ZIKV transmission could leave a considerable amount of naive populations living in regions at risk of ZIKV transmission. Furthermore, expansion of the *Aedes* mosquito distribution, human migration, and shifts in socioeconomic status could lead to more populations being at risk for local ZIKV transmission. It is more likely that ZIKV transmission activities in the future may resemble the current situations of dengue and chikungunya, where transmission activities could flare up sporadically. The possible sporadic outbreaks of ZIKV would continue to pose a risk to the continental US, where most of the population is naive to the virus and a large fraction live in areas environmentally suitable for ZIKV transmission.

## Conclusion

In this study, we show that the overall risk of ZIKV introduction and local transmission is jointly determined by the intensity of ZIKV importations, environmental suitability for ZIKV transmissions and the socioeconomic risk of exposure to mosquitoes. Our estimates suggest that the risk of ZIKV introductions has a very strong spatial and temporal heterogeneity. The areas in the continental US at non-negligible risk (that is, greater than 1/8) only account for 2.56% of the total population in the continental US. The model is able to identify the hotspots for ZIKV introductions, and it reveals the relative contributions of ZIKV introductions from different geographical regions over time. The results of our study have the potential to guide the development of ZIKV prevention and response strategies in the continental US.

## Abbreviations

CDC: Centers for Disease Control and Prevention of the United States
US: United States
TCC: Travel-related Case Counts
ZIKV: Zika virus
GDP: Gross domestic product
PPP: Purchasing power parity

## Author’s contributions

K.S, M.E.H., I.M.L., and A.V. designed research; Q.Z., K.S., M.C., A.P.y.P., S.M., D.M., P.P., L.R., and A.V. performed research; Q.Z., K.S., M.C., A.P.y.P., D.M., and A.V. analyzed data; and Q.Z., K.S., M.C., A.P.y.P., N.E.D., D.P.R., S.M., D.M., P.P., L.R., M.E.H., I.M.L., and A.V. wrote the paper.

